# Excitation spectral microscopy for highly multiplexed fluorescence imaging and quantitative biosensing

**DOI:** 10.1101/2021.04.13.439576

**Authors:** Kun Chen, Rui Yan, Limin Xiang, Ke Xu

## Abstract

The multiplexing capability of fluorescence microscopy is severely limited by the broad fluorescence spectral width. Spectral imaging offers potential solutions, yet typical approaches to disperse the local emission spectra notably impede the attainable throughput. Here we show that using a single, fixed fluorescence emission detection band, through frame-synchronized fast scanning of the excitation wavelength from a white lamp *via* an acousto-optic tunable filter (AOTF), up to 6 subcellular targets, labeled by common fluorophores of substantial spectral overlap, can be simultaneously imaged in live cells with low (∼1%) crosstalks and high temporal resolutions (down to ∼10 ms). The demonstrated capability to quantify the abundances of different fluorophores in the same sample through unmixing the excitation spectra next enables us to devise novel, quantitative imaging schemes for both bi-state and FRET (Förster resonance energy transfer) fluorescent biosensors in live cells. We thus achieve high sensitivities and spatiotemporal resolutions in quantifying the mitochondrial matrix pH and intracellular macromolecular crowding, and further demonstrate, for the first time, the multiplexing of absolute pH imaging with three additional target organelles/proteins to elucidate the complex, Parkin-mediated mitophagy pathway. Together, excitation spectral microscopy provides exceptional opportunities for highly multiplexed fluorescence imaging. The prospect of acquiring fast spectral images without the need for fluorescence dispersion or care for the spectral response of the detector offers tremendous potential.

## Introduction

The wide popularity of fluorescence microscopy in biological research^1,2^ benefits greatly from its two distinct advantages, namely, target specificity and compatibility with live cells. Yet, both advantages connect to performance limits. For multi-target imaging, owing to the broad spectral width of molecular fluorescence, common approaches based on bandpass filters only accommodate 3-4 spectrally well-separated channels within the visible range. Meanwhile, the urge to image fast dynamics in live cells for multiple targets is often impeded by the slow mechanical switching between filter sets. Quantitative imaging of fluorescent biosensors^3-6^, for which the simultaneous monitoring of two or more spectral channels is often necessary, creates yet another level of challenge for fast imaging, and its integration with additional fluorescent markers is even more difficult.

Spectral imaging^7-9^ offers potential solutions to the above problems. Although it is conceptually intuitive to detect the fluorescence emission spectrum, in practice, with common pixel array-based cameras, it remains difficult to record spatial and spectral information at the same time for the wide-field^10,11^. Consequently, spectrally resolved images are usually acquired in a point-scanning manner by dispersing the fluorescence from one single spot of the sample at a time. Alternatively, sparse single molecules may be spectrally dispersed in the wide-field to accumulate super-resolution images over many camera frames^12,13^. Both scenarios place substantial constraints on temporal resolution. Tunable bandpass filters provide a possibility to scan through the emission wavelength in the wide-field^14^. However, applying narrow bandpasses to the fluorescence emission results in inefficient use of the scarce signal. In addition, tunable thin-film filters are limited in speed with mechanical scanning^15^, whereas electronically tunable filters, *e*.*g*., acousto-optic tunable filter (AOTF), only work for polarized light and introduce distortion to wide-field images^16,17^.

In contrast to the above challenges to resolve the fluorescence emission spectrum at every pixel, it is straightforward to scan the excitation wavelength for the entire imaging field while recording the resultant emission in a fixed passband, hence excitation wavelength-resolved images. Excitation spectral microscopy, however, is conceptually less intuitive and experimentally rarely explored^7,18-21^, especially for examining subcellular structural dynamics in live cells. A recent lattice light-sheet setup has imaged 6 subcellular targets using 6 excitation lasers that span the visible range^21^. However, even with large (∼40 nm) spectral separations between the laser lines and between the fluorophore spectra, high (∼50%) target-to-target crosstalks are observed^21^. The implementation of 6 lasers with associated notch filters is also technically demanding.

Here we show that using a standard epifluorescence microscope with a single emission band, through frame-synchronized fast scanning of the excitation wavelength from a white lamp at ∼10 nm resolution, 6 subcellular targets, labeled by common fluorophores of substantial spectral overlap, can be simultaneously imaged in live cells in the wide-field with low crosstalks and high spatiotemporal resolutions. The ability to unmix and quantify different fluorescent species in the same sample *via* the excitation spectrum next enables us to devise fast, quantitative imaging schemes for different modes of fluorescent biosensors in live cells, as well as their multiplexing with multiple additional fluorescent tags.

## Results

### Frame-synchronized AOTF scanning of the excitation wavelength

Light from a white lamp was collimated and linearly polarized before entering an AOTF (Fig. 1a). The 1^st^ order diffracted beam, with its wavelength and intensity controlled by a radio-frequency (RF) synthesizer and polarization orientation rotated by 90° (Ref 17), was cleaned up with a second polarizer and then coupled into a commercial epifluorescence microscope as the excitation source using a conventional, single-band filter cube. Wide-field images collected from a high-NA oil-immersion objective were continuously recorded using an sCMOS or EM-CCD camera. A multifunction I/O device synchronized the RF synthesizer, so that for each successive frame, the excitation switched between 8 preset wavelength profiles that were ∼10 nm apart from each other. The 8 profiles (Supplementary Fig. S1) each had a bandwidth of 5-12 nm, and typical intensities were ∼6 and ∼27 µW for recording at 10 and >200 frames per second (fps), respectively. Full-frame excitation-spectral images were thus obtained every 8 camera frames (Fig. 1b).

**Fig. 1.**
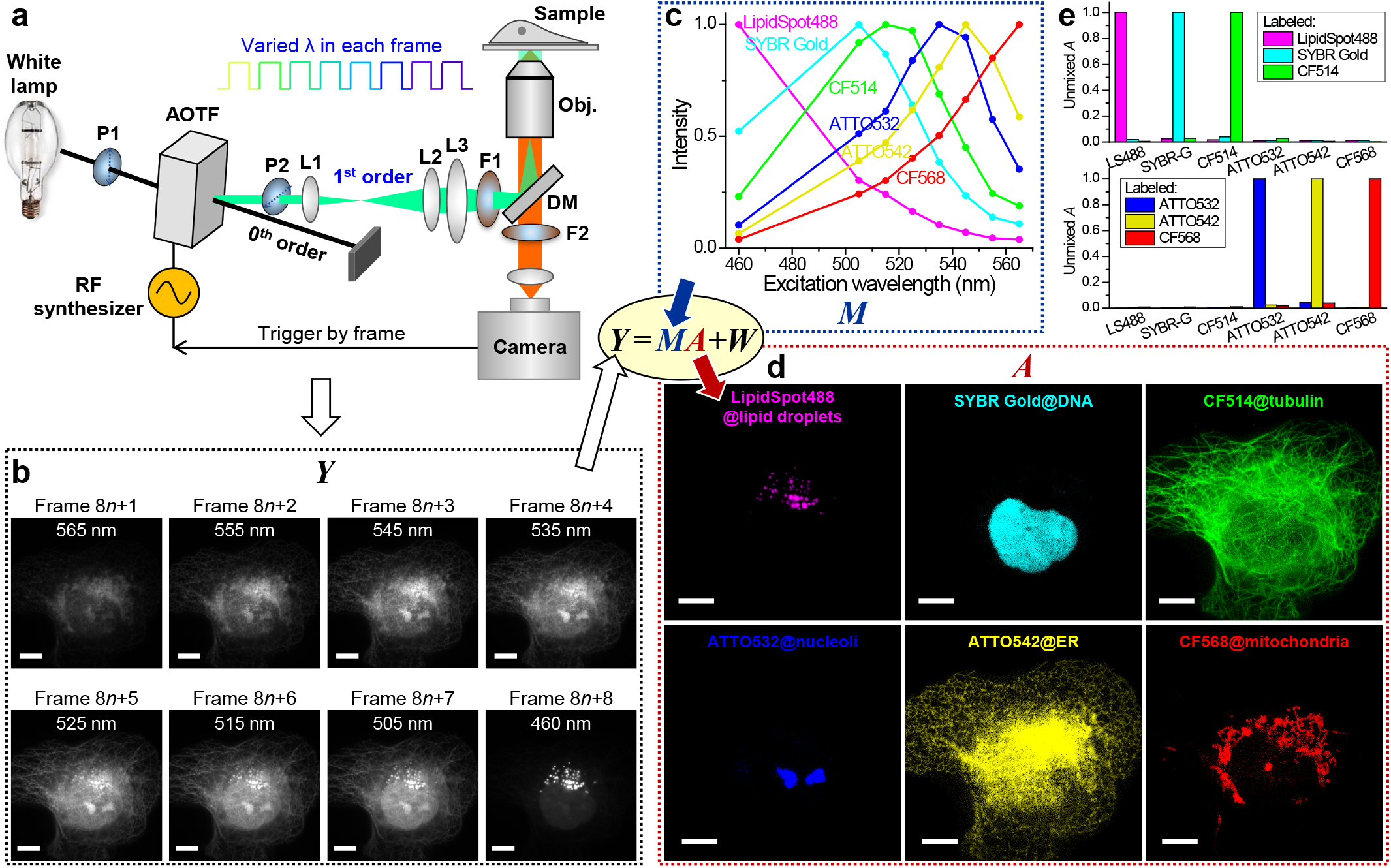
Excitation spectral microscopy. (a) Schematic of the setup. Full-frame spectral micrographs are obtained by the synchronized fast modulation of the excitation wavelength λ in consecutive frames. P, polarizer; L, lens; F, bandpass filter; DM, dichroic mirror. The resultant excitation spectrum collected at every pixel (*Y*) is linearly unmixed into the abundances (*A*) of different fluorophores based on their pre-calibrated excitation spectrum (*M*) to minimize the residual (*W*). (b) Example images recorded at 8 preset excitation wavelengths in 8 consecutive frames at 10 fps (thus 0.8 s total data acquisition time), for a fixed COS-7 cell labeled by 6 fluorescent dyes for 6 distinct subcellular structures: LipidSpot 488 for lipid droplets, SYBR Gold for nuclear DNA, and CF514, ATTO 532, ATTO 542, and CF568 for immunofluorescence of tubulin, nucleoli, ER, and mitochondria, respectively. (c) Reference 8-wavelength excitation spectra of the 6 fluorophores, separately measured on our setup using singly labeled samples. (d) Decomposed images of the 6 fluorophores, obtained via linearly unmixing the excitation-dependent intensity at each pixel in (b) using the reference spectra in (c). (e) Unmixed abundancy values in different fluorophore channels for samples singly labeled by each of the 6 fluorophores. Scale bars: 10 µm (b, d).

### Unmixing 6 fluorophores through the excitation spectrum

As each fluorophore is characterized by its own excitation spectrum, *i*.*e*., a fixed profile of how the fluorescence intensity varies as a function of the excitation wavelength (Fig. 1c), a sample region containing multiple fluorophores should exhibit an excitation spectrum that is a linear combination of that of its component fluorophores^7^. Thus, by linearly unmixing the above-recorded excitation spectrum of every pixel (*Y*) based on the excitation spectrum of each fluorophore (*M*) pre-calibrated on the same setup using singly labeled samples (Fig. 1c), we quantified the local abundance (*A*) of each fluorophore. Rendering the resultant abundance in each pixel as an image thus yielded fluorophore-decomposed micrographs of the sample (Fig. 1d).

Figure 1d shows the unmixed images of a fixed COS-7 cell, in which 6 distinct subcellular structures were labeled with 6 fluorescent dyes that overlapped substantially in the excitation spectrum (Fig. 1c). Little crosstalk was observed between the 6 unmixed images (Fig. 1d) with our 8-wavelength excitation scheme accomplished in 0.8 s (8 frames at 10 fps), even as the raw images at each wavelength (Fig. 1b) contained substantial contributions from multiple fluorophores due to heavy spectral overlaps (Fig. 1c). Forcing the same 6-fluorophore unmixing procedure onto samples singly labeled by each fluorophore showed an average crosstalk of 1.3% between fluorophores (Fig. 1e). Simulations further showed that despite the non-ideal spectral profile of the AOTF output, including side lobes that broaden the spectral widths at the baseline (Supplementary Figs. S1)^17^, low crosstalks were achievable at moderate noise levels for these 6 fluorophores and for the 6 live-cell fluorophores below (Supplementary Figs. S2 and S3).

### Fast multi-target imaging of live cells

We next applied 8 excitation-wavelength spectral microscopy to live cells. With a combination of fluorescent protein (FP) tags and live-cell stains, 6 distinct subcellular targets were well unmixed (Fig. 2a-c) at an averaged 1.3% target-to-target crosstalk (Supplementary Fig. S2). At a moderate camera framerate of 10 fps, a full-frame excitation-spectral image (and thus unmixed 6-target image) was obtained every 0.8 s (Supplementary Video 1). We thus visualized that in live COS-7 cells, lysosome and endoplasmic reticulum (ER) both marked the fission site of a mitochondrion (white arrows in Fig. 2d; Supplementary Video 2), and that the mitochondrial DNA was also divided in this process, consistent with previous, separate observations^22-24^.

**Fig. 2.**
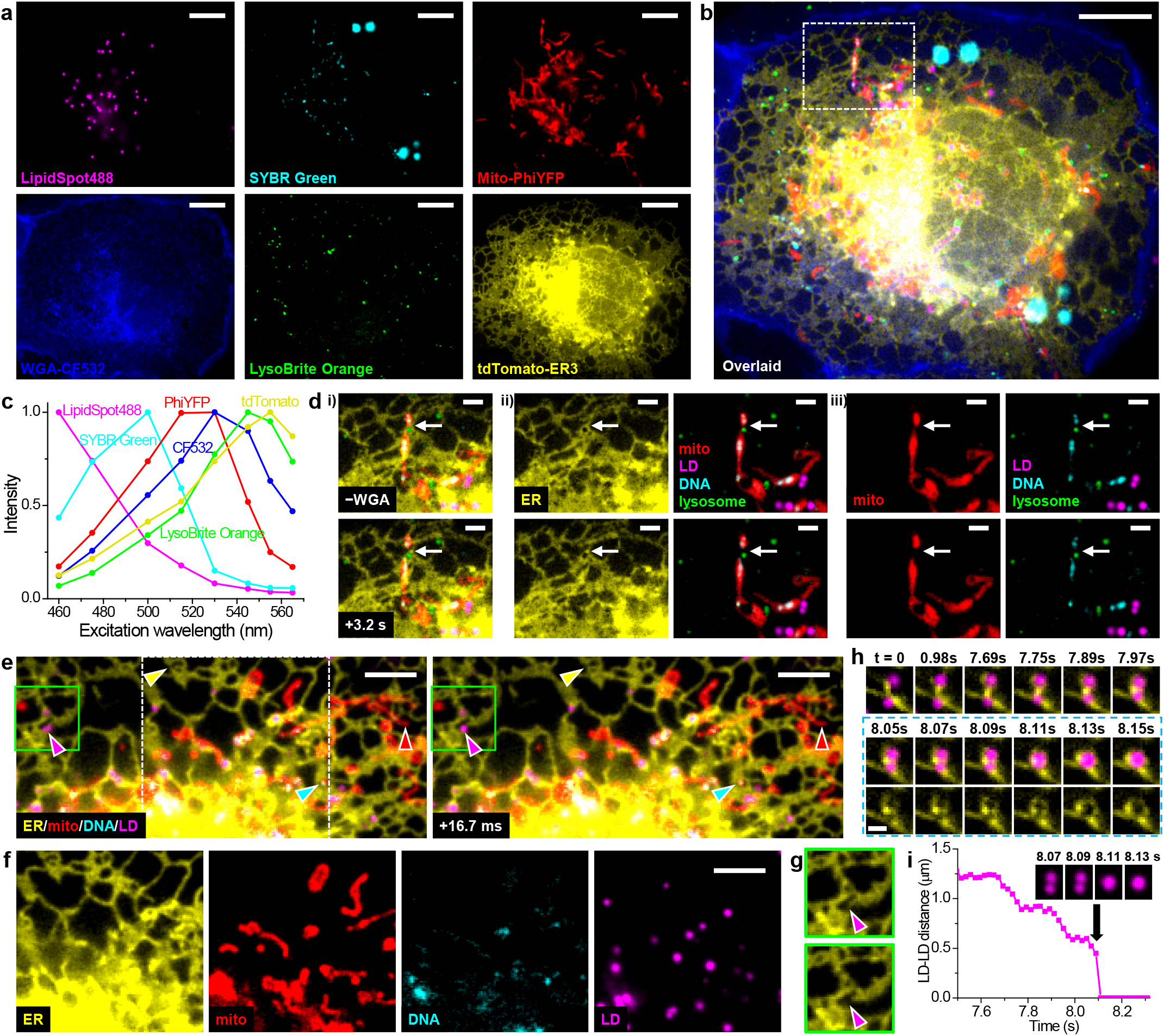
Fast multi-target imaging of live cells. (a) Unmixed images of 6 subcellular targets in a live COS-7 cell via 8-excitation-wavelength recording at 10 fps (0.8 s total data acquisition time). LipidSpot 488: lipid droplets (LDs), SYBR Green: mitochondrial DNA, Mito-PhiYFP: mitochondrial matrix, WGA-CF532: cell membrane, LysoBrite Orange: lysosomes, tdTomato-ER3: ER. (b) Overlay of the above 6 images. (c) Reference excitation spectra of the 6 fluorophores, separately measured on the setup using singly labeled samples. (d) The box region in (b) for two time points 3.2 s apart (top vs. bottom rows), after i) removing the WGA channel, ii) separation of the ER channel, and iii) further separation of the mitochondria channel. (e) Two consecutive 4-fluorophore images at 16.7 ms time spacing for another live COS-7 cell labeled by tdTomato-ER3, Mito-PhiYFP, SYBR Green, and LipidSpot 488, achieved by synchronizing 4-wavelength excitation with 240 fps recording. Yellow, red, cyan, and magenta arrowheads point to noticeable structural changes in each fluorophore channel. (f) The separated fluorophore channels for the white box in (e). (g) The ER channel for the green boxes in (e), at 16.7 ms separation. (h) ER-mediated LD fusion observed in another 4-wavelength experiment at 19.8 ms time spacing (202 fps recording). Bottom row: consecutive ER-LD and ER images during merging. (i) Distance between the two LDs as a function of time. Inset: four consecutive images of the LD channel. Scale bars: 10 µm (a, b); 2 µm (d); 5 µm (e, f); 1 µm (h). See also Supplementary Videos 1-4.

For faster imaging, we next synchronized 4-wavelength excitation cycles with a 240 fps camera framerate (Methods) to image 4 subcellular targets at 16.7 ms time spacing (Fig. 2ef and Supplementary Video 3). This enabled us to simultaneously detect the fast structural changes of ER, mitochondria, mitochondrial DNA, and lipid droplets in the same field of view (color arrowheads in Fig. 2e). In particular, the magenta arrowheads point to the fast fusion of two lipid droplets that occurred within the 16.7 ms timeframe, for which event we also observed associated fast retraction of ER (Fig. 2g). In another 4-wavelength experiment performed at a 202 fps framerate (19.8 ms per spectral image), we recorded how an ER tubule first brought two lipid droplets closer in discrete, ∼0.1 s steps, and then mediated their fusion through fast ER reorganization (Fig. 2hi and Supplementary Video 4), a process substantially different from existing models^25^. Together, we have demonstrated, for both live and fixed cells, the power of excitation spectral microscopy for high-throughput multi-target imaging.

### Absolute pH imaging in live cells via unmixing and quantifying two fluorescent species

We next harnessed our above capability to quantify spectrally overlapped fluorescent species to devise quantitative imaging schemes for fluorescent biosensors in live cells. A class of biosensors works on contrasting fluorescence properties between their analyte-bound and unbound states. The FP pHRed is such a bi-state sensor that exhibits distinct excitation spectra upon protonation and deprotonation^26^. Although this property has enabled ratiometric pH detection with two excitation wavelengths^26,27^, absolute pH mapping is usually not performed.

We achieved absolute pH mapping by quantifying the respective abundances of the protonated (HA) and deprotonated (A^−^) species of pHRed at each pixel. To this end, we first recorded the excitation spectra of pHRed in proton-permeabilized cells at low (4.5) and high (11) pH extremes (Fig. 3a, dash lines), under which conditions the FP should be in the pure HA and A^−^ states, respectively. We next similarly recorded excitation spectra in a series of buffers of pH = 6.5-9.5 (Fig. 3a, solid lines). For each pH, the measured excitation spectrum (e.g., Fig. 3b, solid line) was unmixed into that of the HA and A^−^ states (Fig. 3b, dash lines). The resultant abundances at each pH (Fig. 3c) fitted well to the simple Henderson-Hasselbalch equation based on thermodynamic equilibrium:

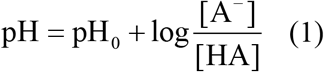

in which pH_0_ is the only fitting parameter, and [A^−^] and [HA] are the abundances of the two states. From the fit we obtained pH_0_ = 7.87, close to the apparent pK_a_ of pHRed (∼7.8) (Ref 26).

**Fig. 3.**
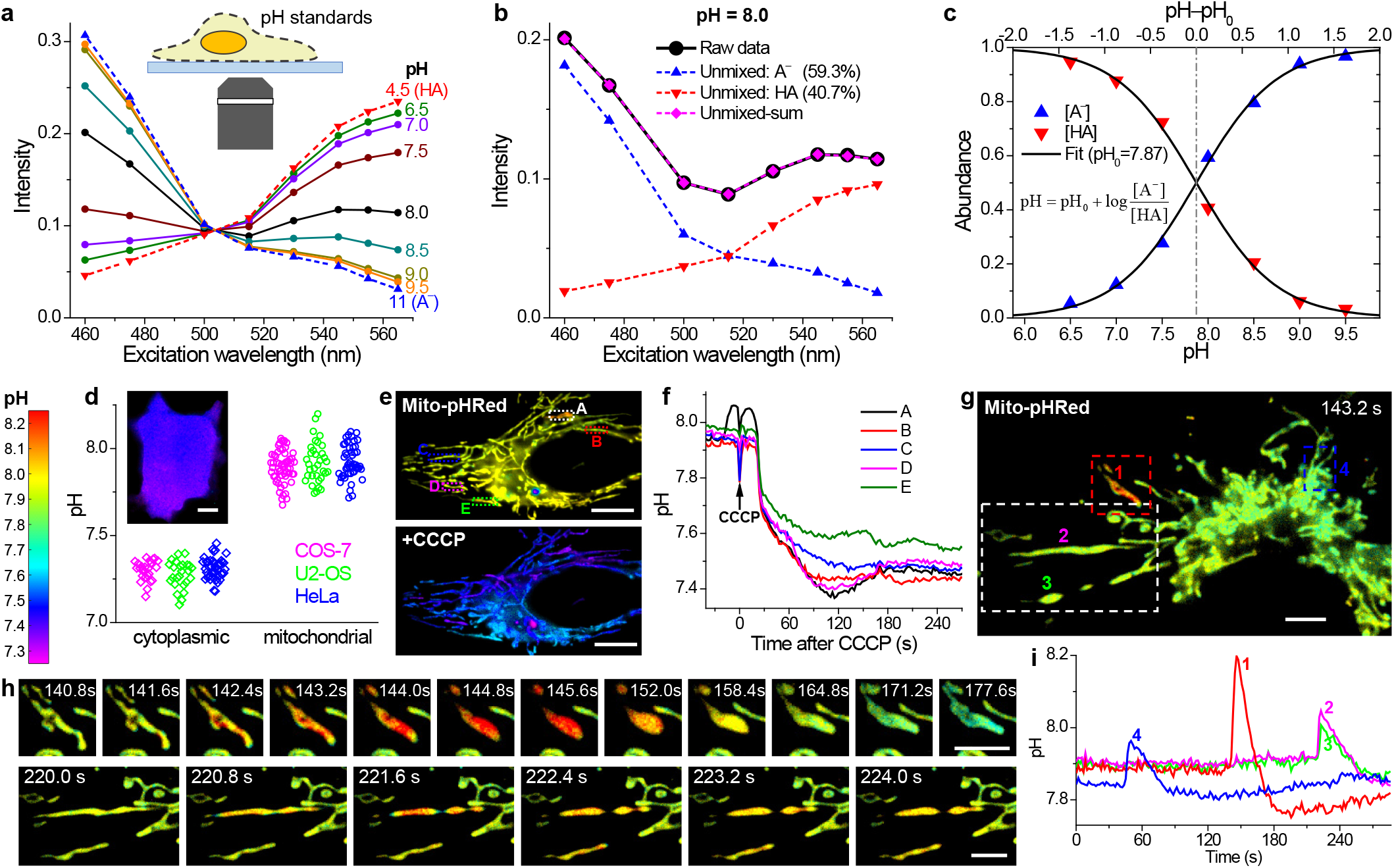
Absolute pH imaging in live cells via unmixing and quantifying two fluorescent species. (a) Excitation spectra measured by our spectral microscope, for pHRed in proton-permeabilized COS-7 cells in buffer standards of pH = 4.5 and 11 (dash lines; taken as pure forms of HA and A^−^, respectively), and of pH = 6.5-9.5 (solid lines). (b) Linear unmixing of the excitation spectrum at pH = 8.0 (black solid line) into HA and A^−^ components (dash lines). (c) Fitting the resultant HA and A^−^ abundances unmixed at different pH values to the Henderson-Hasselbalch equation with a single parameter pH_0_. (d) Statistics of the average pH values of the cytoplasm and mitochondrial matrix of live COS-7, U2-OS, and HeLa cells, measured with our approach using cytoplasmic pHRed and Mito-pHRed, respectively. Inset: Color-coded absolute pH map of the cytoplasm of a live COS-7 cell. (e) Mito-pHRed absolute pH maps [same color scale as (d)] of the mitochondrial matrix in a live HeLa cell, before (top) and after (bottom) 120 s treatment with 20 µM CCCP. (f) pH value time traces for the five regions marked in (e). (g) Color-coded Mito-pHRed absolute pH map of the mitochondrial matrix in a live COS-7 cell, at the time point of 143.2 s. (h) Time sequences for the red and white boxed regions in (g), for two different time windows. (i) pH value time traces for the four mitochondria marked in (g). Experiments were performed with 8-wavelength excitation cycles at 10 fps, corresponding to 0.8 s acquisition time for each spectral image. Scale bars: 10 µm (d, e); 5 µm (g, h). See also Supplementary Videos 5-7.

This good fit to theory allowed us to image absolute pH in live cells by first unmixing the excitation spectrum at every pixel and then feeding the resultant HA and A^−^ abundances into eqn 1. Besides achieving high confidences for HA and A^−^ abundances and hence pH, this approach enabled us to readily incorporate additional fluorophores, as discussed below. For pHRed expressed in the cytoplasm, we observed homogenous pH of ∼7.2-7.4 in common cell lines (Fig. 3d). Meanwhile, Mito-pHRed in the mitochondrial matrix showed typical pH of ∼7.7-8.1 (Fig. 3d-g) in live cells while exhibiting sporadic, spontaneous jumps (Fig. 3hi). Treating the cells with the uncoupler carbonyl cyanide *m*-chlorophenyl hydrazone (CCCP) led to fast drops of the mitochondrial matrix pH to ∼7.4-7.5 (Fig. 3ef). While these results are qualitatively consistent with previous observations^26-28^, our full-frame fast recording quantified absolute pH at high spatiotemporal resolutions. For the CCCP-treated cells, we thus found diverse pH dynamics between different mitochondria in the same cell (Fig. 3f and Supplementary Video 5). For the spontaneous pH jumps in untreated live cells, we showed that the matrix pH of individual mitochondria abruptly rose by ∼0.1-0.3 units in ∼2 s, and such fast processes often coincided with abrupt mitochondrial shape changes that were fittingly captured with our 0.8 s full-frame time resolution (Fig. 3hi, Supplementary Figs. S4 and S5, and Supplementary Videos 6 and 7). The pH then gradually decreased in ∼30 s and overshot to a value ∼0.05 lower than the starting pH, a process often accompanied by further changes in mitochondrial shape (Fig. 3hi, Supplementary Fig. S4, and Supplementary Videos 6 and 7).

### FRET imaging in live cells via resolving the excitation spectrum

We next examined another major class of biosensors based on Förster resonance energy transfer (FRET). Although modern FRET imaging often focuses on separating the fluorescence emissions of the donor and acceptor fluorophores^4,6^, early FRET spectroscopy studies favored the analysis of the excitation spectrum^29^: when monitoring the emission from the acceptor, FRET gives rise to new features in the excitation spectrum corresponding to donor absorption.

We achieved a microscopy version of this spectroscopy approach using the Clover-mRuby2 FP pair^30^. As our emission band well-matched the emission of mRuby2 but not Clover, for COS-7 cells co-expressing the two free FPs, the recorded 8-wavelength excitation spectrum was dominated by mRuby2 (Fig. 4a, solid line). Unmixing this spectrum into mRuby2 and Clover components (Fig. 4a, dash lines) showed the peak height of the latter to be ∼7% of the former. In contrast, for a construct in which Clover and mRuby2 were directly linked, the excitation spectrum showed a substantial rise at ∼500 nm (Fig. 4b, solid line), attributable to FRET-induced mRuby2 emission due to Clover absorption^29^. Unmixing the spectrum into mRuby2 and Clover components (Fig. 4b, dash lines) showed the peak height of the latter to be ∼62% of the former. Subtracting the above ∼7% bleed-through from Clover emission thus yielded a ∼0.55 ratio between the peak heights due to the FRET donor and acceptor. Given the near-identical peak extinction coefficients of mRuby2 and Clover^30^, this ∼0.55 ratio may be taken as the FRET transfer efficiency *E* (Methods)^29^, in agreement with emission spectroscopy results^30^ on a similar construct (0.55).

**Fig. 4.**
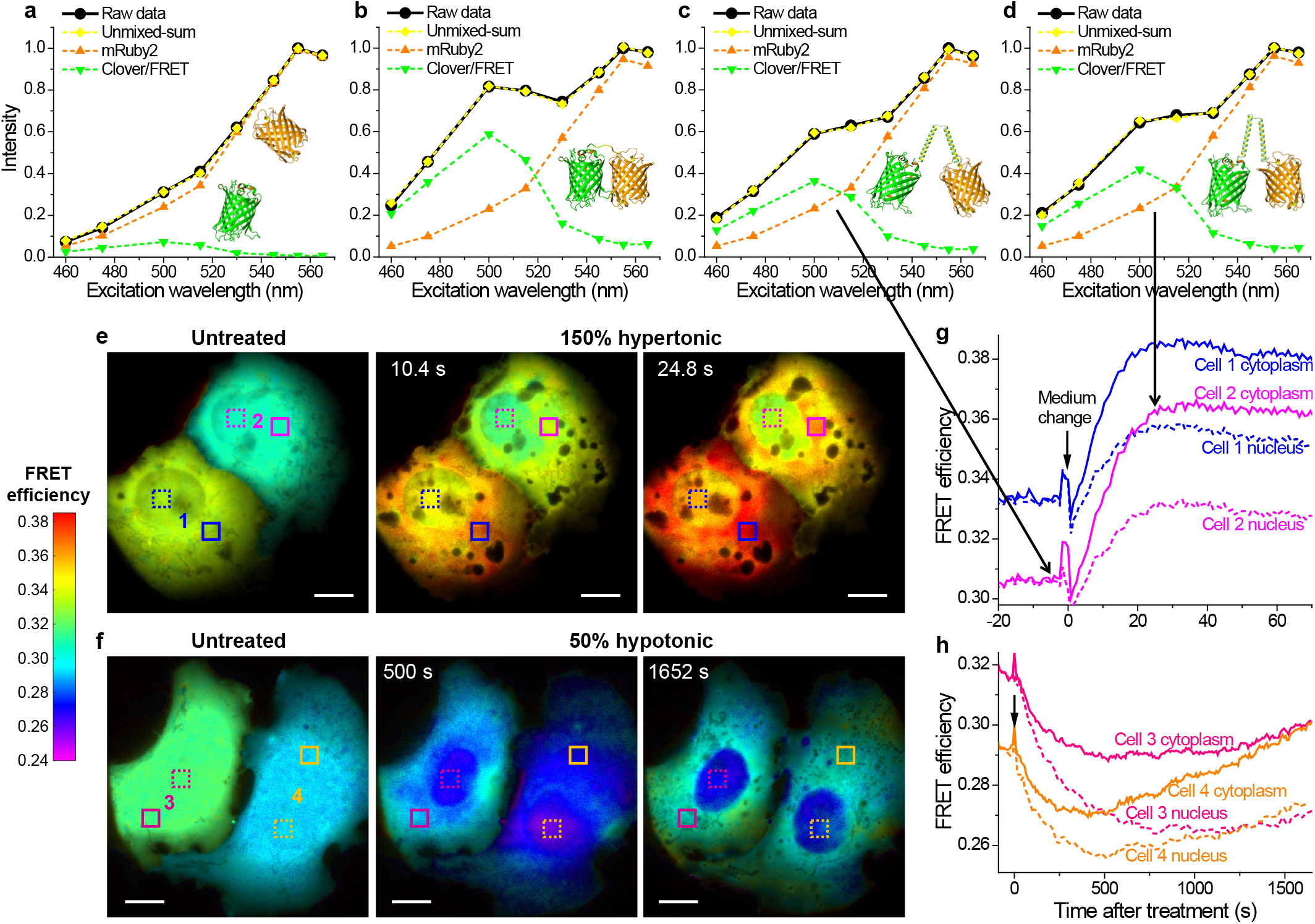
FRET imaging in live cells via resolving the excitation spectrum. (a) Excitation spectrum measured by our spectral microscope (black solid line) and its unmixing (dash lines) for non-interacting mRuby2 and Clover co-expressed in the cytoplasm of a live COS-7 cell. (b) Measured excitation spectrum and its unmixing for a directly linked Clover-mRuby2 construct expressed in a live COS-7 cell. (c,d) Measured excitation spectrum and its unmixing for the Clover-mRuby2 FRET crowding sensor in the cytoplasm of a live COS-7 cell, before (c) and ∼25 s after (d) 150% hypertonic treatment by adding into the cell medium an equal volume of medium that was supplemented with 300 mM sorbitol. (e) Color-coded FRET maps for the crowding sensor, for two live COS-7 cells before (left), ∼10 s after (center), and ∼25 s after (right) the 150% hypertonic treatment. (f) Color-coded FRET maps for the crowding sensor, for two live COS-7 cells before (left), 500 s after (center), and 1650 s after (right) 50% hypotonic treatment by adding into the medium an equal volume of water. (g, h) FRET value time traces for the boxed regions of cytoplasm (solid lines) and nuclei (dash lines) of the 4 cells in (e, f). Experiments were performed with 8-wavelength excitation cycles at 10 fps (0.8 s acquisition time for each spectral image). Scale bars: 10 µm (e, f). See also Supplementary Videos 8-9.

We next constructed a FRET sensor for quantifying macromolecular crowding in live cells by linking Clover and mRuby2 with a long, conformationally flexible domain^31^. When expressed in the COS-7 cell cytoplasm, this sensor exhibited lower FRET efficiencies (Fig. 4cd) than the directly linked Clover-mRuby2 (Fig. 4b), consistent with increased donor-acceptor distances. Upon treating the live cells with 150% and 50% osmotic pressures of the normal medium, the measured FRET efficiency rose and dropped substantially (Fig. 4d-h), as expected for the reduced and increased donor-acceptor distances under raised and lowered levels of macromolecular crowding inside the cells, respectively^31^.

Through continuous recording under our 8-wavelength excitation scheme at 10 fps and performing the above FRET analysis for every pixel, we obtained full-frame FRET images at 0.8 s time resolution, and thus unveiled rich spatiotemporal heterogeneities in macromolecular crowding in live cells (Fig. 4ef). In untreated COS-7 cells, the observed FRET efficiency was largely uniform within each cell, yet varied substantially (∼0.29-0.33) between cells. Interestingly, whereas the 150% hypertonic treatment resulted in quick rises in the intracellular FRET signal that plateaued in ∼25 s (Fig. 4e and g and Supplementary Video 8), the 50% hypotonic treatment led to slow decreases in the FRET signal over ∼500 s and subsequent recoveries (Fig. 4fh and Supplementary Video 9). These results may be rationalized with the contrasting, quick reduction versus slow expansion-recovery of cell volumes under hypertonic and hypotonic conditions^32,33^. At the subcellular level, we found that for the hypertonic treatment, the nuclear FRET signals rose slower and achieved lower final increases when compared to the cytoplasm (∼0.03 vs. ∼0.05) (Fig. 4eg). This result may indicate that the nuclear envelop retarded volume reduction and thus the rise in macromolecular crowding. In contrast, for the opposite, slow hypotonic process, the cytoplasmic FRET signals dropped slower than the nuclei, with some cells further showing complete recovery over long periods of time (Fig. 4fh). This result may be attributed to the regulatory volume decrease processes in the cytoplasm^34^.

### Combining absolute pH imaging with three additional FP tags to elucidate the Parkin-mediated mitophagy pathway

We conclude by multiplexing our above quantitative biosensing and multi-fluorophore imaging capabilities. To probe a biologically interesting multistate system, we examined the Parkin-mediated mitophagy pathway^35,36^. For live HeLa cells lacking Parkin, absolute pH imaging with Mito-pHRed showed that after ∼30 min CCCP treatment, the mitochondrial matrix pH gradually recovered from the initial drop to reach ∼7.7 at ∼1 h (Fig. 5ac and Supplementary Figs. S6 and S7). Relatively homogenous pH values were detected between different mitochondria in the same cell, yet some cell-to-cell variations were noticed. Remarkably, under the same experimental conditions, HeLa cells expressing fluorescently untagged Parkin exhibited a large heterogeneity in mitochondrial matrix pH (∼6.7-7.7), with notably dissimilar pH values observed for neighboring mitochondria in the same cell (Fig. 5bc and Supplementary Fig. S7).

**Fig. 5.**
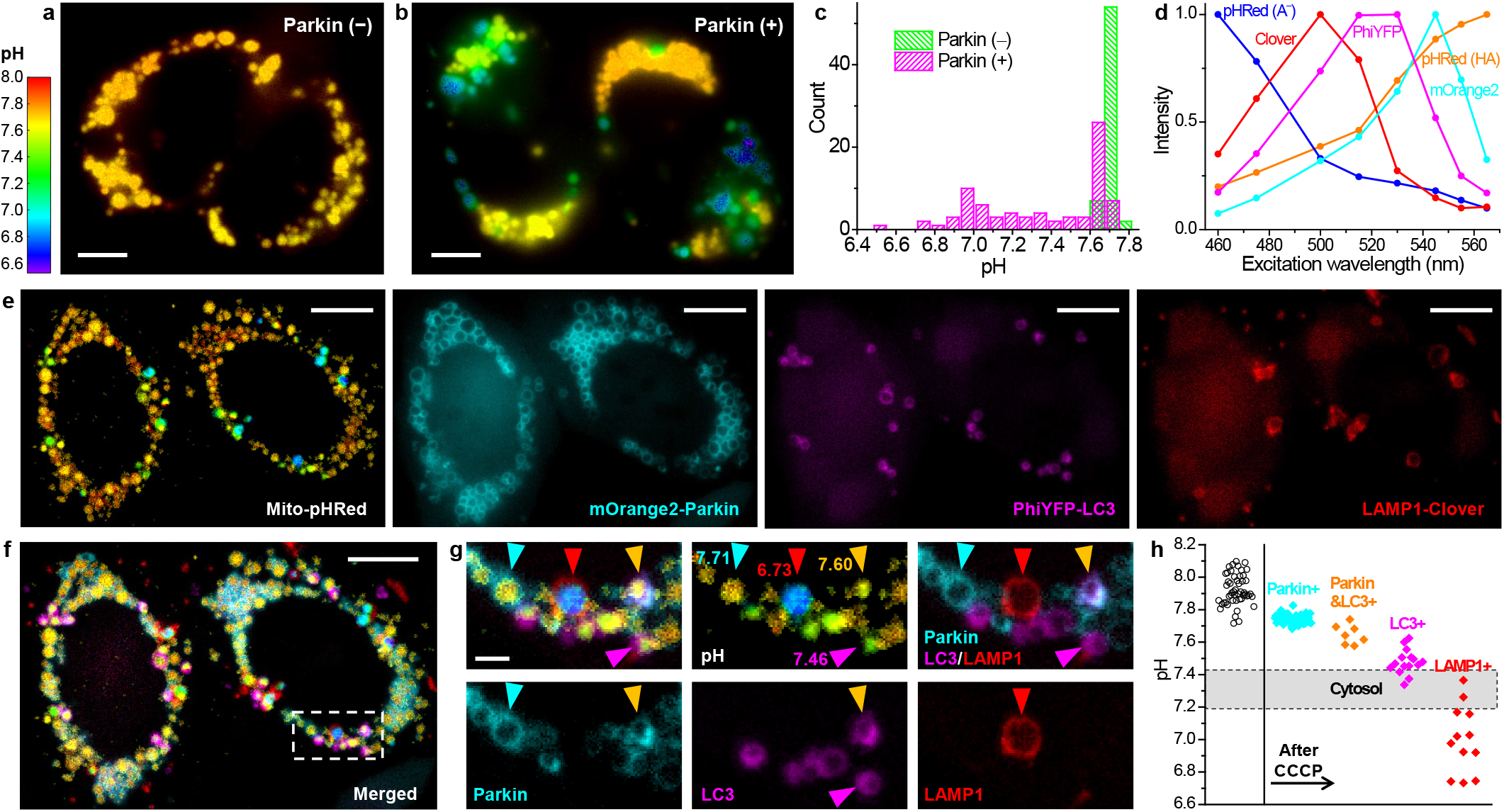
Concurrent absolute pH imaging of the mitochondrial matrix with three additional FP-tagged markers in the Parkin-mediated mitophagy pathway. (a, b) Color-coded Mito-pHRed absolute pH maps of live HeLa cells after the application of 20 µM CCCP for 6 h, for cells without (a) and with (b) the co-expression of fluorescently untagged Parkin. (c) Distribution of the measured pH values in each mitochondrion for (a) and (b). (d) Excitation spectra on our setup for the five fluorescent species unmixed to enable absolute pH imaging with three additional FP markers. (e) Unmixed images of color-coded Mito-pHRed absolute pH map, mOrange2-Parkin, PhiYFP-LC3, and LAMP1-Clover for two Parkin-expressing live HeLa cells after the application of 20 µM CCCP for 4 h. (f) Overlay of the images in (e). (g) Zoom-in of the white box in (f), as well as its separation into different channels. Cyan, magenta, and red arrowheads point to individual mitochondria only marked by Parkin, LC3, and LAMP1, with respective matrix pH values quantified from the enclosed Mito-pHRed signals indicated. Orange arrowhead points to a mitochondrion marked by both Parkin and LC3. (h) Distribution of the Mito-pHRed-quantified matrix pH of individual mitochondria in the sample labeled by the different markers (color diamonds), compared to that in untreated HeLa cells (black circles). Shaded band: typical range of cytoplasmic pH. Experiments were performed with 8-wavelength excitation at 10 fps (0.8 s acquisition time for each spectral image). Scale bars: 5 µm (a, b); 10 µm (e, f); 2 µm (g).

To examine whether these different mitochondrial pH states could correspond to different stages of mitophagy^35,36^, we next co-expressed three FP-tagged mitophagy-stage markers, mOrange2-Parkin, PhiYFP-LC3, and LAMP1-Clover, in the cell and performed similar long-term CCCP treatments. The five fluorescent species (A^−^ and HA forms of pHRed, plus mOrange2, PhiYFP, and Clover; Fig. 5d) were well unmixed with our 8-wavelength excitation spectral microscopy (Fig. 5e-g). Scrutiny of the Mito-pHRed-quantified pH values at mitochondria enclosed by the three FP-tagged markers (*e*.*g*., Fig. 5g) showed that the Parkin-marked mitochondria, at the onset of mitophagy, had matrix pH of ∼7.7-7.8 (cyan arrowheads in Fig. 5g; Fig. 5h), comparable to the high-pH population in the Parkin-expressing cells without FP-tagged markers (Fig. 5b and Supplementary Fig. S7). Meanwhile, mitochondria positive for both Parkin and the autophagosome marker LC3 exhibited lower matrix pH values of ∼7.5-7.7 (orange arrowheads in Fig. 5g; Fig. 5h). Mitochondria marked by LC3 alone showed yet lower matrix pHs of ∼7.3-7.6 (magenta arrowhead in Fig. 5g; Fig. 5h). The lower values in this distribution matched cytoplasmic pH (shaded band in Fig. 5h), suggesting complete loss of mitochondrial function. Lastly, mitochondria enclosed by the lysosome marker LAMP1 showed the lowest pHs (<∼7.4) (red arrowhead in Fig. 5g; Fig. 5h), suggesting gradual acidification of the mitochondrial matrix after lysosome fusion. To further validate these results, we permutated the FPs tagged to the above three mitophagy-stage markers. The resultant spectrally resolved images of Mito-pHRed, PhiYFP-Parkin, Clover-LC3, and LAMP1-mOrange2 showed comparable matrix pH trends for mitochondria positive for the different stage markers (Supplementary Fig. S8). We thus elucidated a model in which the mitochondrial matrix pH progressively decreases along the mitophagy pathway. These results contrast with previous efforts employing pH-sensitive FPs to detect mitophagy^37,38^, in which only the mitophagy end results are detected and absolute pH is not measured to elucidate the pathway. Moreover, with the fast full-frame imaging capability, we further showed that the LAMP1-enclosed mitochondria, with their characteristic low matrix pH, moved much faster when compared to their Parkin- and LC3-enclosed counterparts (Supplementary Fig. S8 and Supplementary Video 10), reflecting the high mobility of lysosomes in cells.

## Discussion

In this work, we demonstrated excitation spectral microscopy as a powerful tool for both fast multi-target imaging and quantitative biosensing, as well as their combined use. Whereas previous spectral-imaging efforts often focus on resolving the emission spectrum, the need to disperse the local spectra impedes applications to high-throughput imaging in the wide-field. Meanwhile, as recent experiments probe the dynamics of 6 subcellular targets by exciting with 6 lasers that span the entire visible range, high (∼50%) crosstalks are found in spite of the large spectral separation^21^. Separately, although fluorescence lifetime imaging and stimulated Raman scattering microscopy have recently enabled highly multiplexed imaging^39,40^, dedicated optical setups and/or labeling probes are required, and they cannot be readily applied to fast wide-field imaging.

We showed that under the typical wide-field fluorescence microscopy scheme with a conventional single-band filter cube, by fast scanning the excitation wavelength from a low-cost white lamp, 6 subcellular targets, labeled by common fluorophores of substantial spectral overlap, can be simultaneously imaged in live cells at low (∼1%) crosstalks. By synchronizing the excitation wavelength to each camera frame through fast electronic modulation of the AOTF, full-frame excitation spectral images were obtained with no moving parts, hence high temporal resolutions that were effectively only limited by the camera framerate and the desired number of excitation wavelengths. As an additional benefit, as fluorescence detection was performed for a fixed emission band, the measured spectral behaviors were robustly defined by the preset excitation profiles and independent of the spectral response of the detection system. We thus were able to freely switch between different sCMOS and EM-CCD cameras in this study without the need to modify or recalibrate the system. The potential extension of our approach to even more fluorophores may be achieved by further increasing the number of excitation wavelengths or integrating emission dispersion, although the latter approach would encounter the same throughput limitations we set out to overcome in the first place. Practically, the ultimately achievable multi-target imaging capability is further limited by how well different intracellular targets may be concurrently and specifically labeled.

The capability to unmix and quantify different, spectrally overlapped fluorescent species in the same sample via the excitation spectrum next enabled us to devise fast, quantitative imaging schemes for fluorescent biosensors in live cells, as well as their multiplexing with additional fluorophores. For the bi-state sensor pHRed, we quantified the pH-dependent abundances of its protonated and deprotonated states, and well fitted them to the ideal Henderson-Hasselbalch equation to enable absolute pH imaging, thus visualizing concurrent fast changes in the mitochondrial shape and matrix pH. For FRET imaging, we established a microscopy version of the early excitation spectroscopy approaches to quantify FRET efficiency, and thus achieved high sensitivities and spatiotemporal resolutions in unveiling rich dynamics for macromolecular crowding in live cells. To conclude, we integrated our biosensor and multi-fluorophore imaging capabilities, and achieved absolute pH imaging with three additional FP tags to elucidate the Parkin-mediated mitophagy pathway, thus unveiling fascinating spatiotemporal dynamics for this complex, multistate system.

Together, our results unveil the exceptional opportunities excitation spectral microscopy provides for highly multiplexed fluorescence imaging. Whereas in this work we focused on a facile system based on a lamp-operated epifluorescence microscope, the fast multi-fluorophore and quantitative biosensor imaging capabilities we demonstrated here should be readily extendable to other systems, including light-sheet fluorescence microscopy^2,41^ and structured illumination microscopy^42^, for which cases frame-synchronized fast wavelength scanning may be implemented with supercontinuum light sources^43^. The prospect of acquiring fast spectral images in the wide-field without the need for fluorescence dispersion or the care for the spectral response of the detector offers tremendous potential.

## Materials and methods

### Optical setup

White light from a plasma lamp (HPLS301, Thorlabs) was collimated into a beam ∼4 mm in diameter, and linearly polarized through a wire grid polarizer (WP25M-VIS, Thorlabs). The polarized beam entered an acousto-optic tunable filter (AOTF) (EFLF100L1, Panasonic), and the 1^st^ order diffracted beam, with its polarization direction rotated by 90° versus the incident beam^17^, was cleaned up with a second polarizer (WP25M-VIS) mounted perpendicular to the first polarizer (Fig. 1). This excitation beam was coupled into an Olympus IX73 inverted epifluorescence microscope and focused at the back focal plane of an oil-immersion objective lens (Olympus, UPLSAPO100XO, NA 1.40), thus illuminating a sample area of ∼70 µm diameter. The excitation filter, dichroic mirror, and emission filter used were FF01-505/119, FF573-Di01, and FF01-630/92, respectively, from Semrock. The AOTF unit was driven by an 8-channel RF synthesizer (97-03926-12, Gooch & Housego). The excitation wavelength profiles at different applied RF frequencies (Supplementary Fig. S1) were measured using a visible spectrometer (USB4000-VIS-NIR, Ocean Optics), with power levels determined by a photodiode power sensor (S120VC, Thorlabs). Typical excitation power was ∼6 µW [for recording at 10 frames per second (fps)] and ∼27 µW (for recording at 240 and 202 fps) at each wavelength, corresponding to a power density of ∼0.16-0.72 W/cm^2^ at the sample, which allowed adequate signal and minimal photobleaching. The prolonged recording in Fig. 4fh was performed with lowered excitation powers of ∼2 µW. Wide-field fluorescence images were continuously recorded using an sCMOS camera (Zyla 4.2, Andor) at an effective pixel size of 130 nm (after 2×2 binning) for 512×512 pixels (67×67 µm^2^ field of view) at 10 fps or for 512×150 pixels (67×20 µm^2^ field of view) at 202 fps, or an EMCCD camera (iXon Ultra 897, Andor) at an effective pixel size of 160 nm for 448×110 pixels (72×18 µm^2^ field of view) at 240 fps. The camera was internally triggered, and the “fire” (exposure) output trigger signal was read by a multifunction I/O board (PCI-6733, National Instruments), which in turn modulated the RF synthesizer for the synchronized control of the excitation wavelength in each camera frame (below).

### Spectral imaging via fast AOTF scanning of the excitation wavelength

In most experiments, the 8 channels of the RF synthesizer were preset to 8 different fixed RF frequencies and powers, hence 8 fixed excitation wavelengths and intensities for the AOTF output. These 8 excitations were sequentially applied to each successive camera frame in cycles, so that full-frame excitation-spectral images were obtained every 8 consecutive camera frames. For the 6-target imaging of fixed cells, the preset AOTF driving frequencies were 61.567, 63.400, 65.148, 66.749, 68.454, 70.277, 71.500, and 82.370 MHz, corresponding to excitations centered at 565, 555, 545, 535, 525, 515, 505, and 460 nm, respectively (Supplementary Fig. S1), and they were synchronized with 10 fps sCMOS recording. For the 6-target imaging of live cells and all biosensor experiments, the 8 AOTF driving frequencies were 61.567, 63.400, 65.148, 66.995, 70.277, 72.700, 78.420, and 82.370 MHz, corresponding to excitations centered at 565, 555, 545, 530, 515, 500, 475, and 460 nm, respectively (Supplementary Fig. S1), and they were also synchronized with 10 fps sCMOS recording. For the 4-excitation wavelength imaging of fast dynamics in live cells, 4 RF channels were set to 63.400, 70.277, 72.700, and 82.370 MHz, corresponding to excitations centered at 555, 515, 500, and 460 nm, respectively, and they were synchronized with 240 fps EMCCD or 202 fps sCMOS recordings.

### Acquiring the reference excitation spectra

The reference excitation spectrum of each fluorophore was obtained from singly labeled live and fixed cells by recording images at the above preset excitation wavelengths on our setup. For pHRed, excitation spectra at different pH values (Fig. 3a) were obtained from pHRed-expressing COS-7 cells that were proton-permeabilized with valinomycin and nigericin and immersed in a series of buffer standards (Intracellular pH Calibration Buffer Kit, P35379, ThermoFisher, plus additional homemade buffers).

### Linear unmixing of the experimentally recorded excitation spectra

For both multi-fluorophore and biosensor imaging, the recorded excitation spectrum at each pixel, *e*.*g*., collected from 8 consecutive frames of raw images, was each unmixed into a linear combination of the reference excitation spectra (above) of the pertaining fluorophores using least squares regression under non-negativity constraints^7,8^. The resultant abundances of different fluorophores at each pixel were directly used to generate unmixed images for multi-target imaging or processed further for quantitative biosensor imaging, as described in the text.

### Calculation of the FRET efficiency

The use of excitation spectra to calculate the FRET efficiency is established in earlier studies^29^: for a FRET system consisting of a donor and an acceptor, if one monitors the fluorescence emission of the acceptor, the excitation spectrum *F* is a linear combination of the absorption spectra of the acceptor and the donor: *F* = *ε*_A_ + *Eε*_D_. Here *E* is the FRET efficiency, and *ε*_A_ and *ε*_D_ are the wavelength-dependent extinction coefficients of the acceptor and the donor, respectively. In our experiments, for each pixel we experimentally obtained *F*, and linearly unmixed it into the excitation (absorption) spectra of the donor and the acceptor (above). *E* may thus be determined from the ratio between the peak heights of the donor and acceptor, scaled by the ratio between the known peak extinction coefficients of the acceptor and the donor. For our FRET pair, the peak extinction coefficients Clover and mRuby2 are near identical^30^ (111 vs. 113 mM^−1^ cm^−1^). We thus took the peak height ratios between the donor and acceptor (minus the ∼7% bleed-through from the donor emission) directly as the FRET efficiency.

### Plasmids

tdTomato-ER-3 and LAMP1-Clover (Clover-Lysosomes-20) were gifts from Michael Davidson (Addgene #58097 and #56528). Mito-PhiYFP (pPhi-Yellow-mito) was from Evrogen (#FP607). GW1-pHRed and GW1-Mito-pHRed were gifts from Gary Yellen (Addgene #31473 and #31474) for expression of pHRed in the cytoplasm and mitochondrial matrix, respectively^26^. The Clover-mRuby2 FRET crowding sensor was constructed by inserting a conformationally flexible linker^31^ [(GSG)_6_A(EAAAK)_6_A(GSG)_6_A(EAAAK)_6_A(GSG)_6_; synthesized by Twist Bioscience] between the Clover and mRuby2 of pcDNA3.1-Clover-mRuby2 (Addgene #49089; a gift from Kurt Beam) at the AgeI site. The co-expression plasmid of free Clover and mRuby2 was constructed by inserting an internal ribosome entry site (cloned from Addgene #127332; a gift from Jeremy Wilusz) between the Clover and mRuby2 of pcDNA3.1-Clover-mRuby2 at the AgeI site. For the expression of untagged Parkin, pCMV-Parkin was constructed by inserting Parkin (cloned from Addgene #89299, a gift from Michael Lazarou) into an N1 cloning vector (cut from Addgene #54642, a gift from Michael Davidson). mOrange2-Parkin was constructed by first inserting mCherry-Parkin (cloned from Addgene #59419, a gift from Richard Youle) into an N1 cloning vector (cut from Addgene #54642, a gift from Michael Davidson), and then replacing mCherry with mOrange2 (cloned from Addgene #57969, a gift from Michael Davidson) between the BmtI and BspEI sites. PhiYFP-Parkin was constructed by replacing mOrange2 in the mOrange2-Parkin with PhiYFP between the BmtI and BspEI sites. PhiYFP-LC3 was constructed by replacing mRFP in the pmRFP-LC3 (Addgene #21075; a gift from Tamotsu Yoshimori) with PhiYFP between the BmtI and BglII sites. Clover-LC3 was constructed by replacing mRFP in the pmRFP-LC3 with Clover between the BmtI and BglII sites. LAMP1-mOrange2 was constructed by replacing mRuby2 in the LAMP1-mRuby2 (Addgene #55902; a gift from Michael Davidson) with mOrange2 between the AgeI and NotI sites.

### Live-cell experiments

COS-7, U2-OS, and HeLa cells (Cell Culture Facility, UC-Berkeley) were cultured in Dulbecco’s modified Eagle’s medium (DMEM) (Gibco 31053-028) supplemented with 10% fetal bovine serum (Corning), 1× GlutaMAX Supplement, and 1× non-essential amino acids, at 37 °C and 5% CO_2_. For live-cell experiments, cells were cultured in Lab-Tek 8-well chambered coverglass (ThermoFisher), and transiently transfected with the above plasmids, either alone or in combination. Transfection was performed using Lipofectamine 3000 (ThermoFisher) or the Neon Transfection System (ThermoFisher), following the manufacturers’ instructions. For live-cell staining of the plasma membrane, lipid droplet, and DNA, wheat germ agglutinin (WGA) CF532 (300-500×, 29064, Biotium), LipidSpot488 (300-1000×, 70065, Biotium), and SYBR Green (15000-100000×, S7536, ThermoFisher) were added to the medium for 30 min at 37 °C and washed 3 times with DMEM before imaging. Imaging buffer was the regular culture medium with the addition of 25 mM HEPES at pH 7.4 (15630106, Gibco) or a commercial buffer based on MOPS (Hibernate A, BrainBits), with similar results observed.

### Fixed-cell experiments

For fixed-cell experiments, COS-7 cells were plated in an 8-well LabTek chamber at ∼30% confluency. After 24 h, cells were fixed using 3% paraformaldehyde and 0.1% glutaraldehyde in phosphate-buffered saline (PBS) followed by two washes with 0.1% sodium borohydride in PBS. Cells were blocked and permeabilized in a blocking buffer (3% bovine serum albumin with either 0.5% Triton X-100 or 0.02% saponin in PBS), followed by primary and secondary antibody labeling. Primary antibodies used were chicken anti-α-tubulin (ab89984, Abcam) for labeling of microtubules, rabbit anti-Nogo (ab47085, Abcam) for labeling of ER, mouse IgG1 anti-NPM1 (32-5200, ThermoFisher) for labeling of nucleoli, and mouse IgG2a anti-Tom20 (sc-17764, Santa Cruz Biotech) for labeling of mitochondria. Secondary antibodies (Jackson ImmunoResearch) were labeled via reaction with dye NHS esters: CF514 (92103, Biotium), CF568 (92131, Biotium), ATTO 532 (AD532-31, ATTO-TEC) and ATTO 542 (AD542-31, ATTO-TEC). After antibody staining, cells were incubated in LipidSpot488 (300-1000×, 70065, Biotium) and SYBR Gold (300-1000×, S11494, ThermoFisher) in PBS to stain lipid droplets and nuclear DNA for 30 min, and the sample was washed with PBS for 10 min × 3 times before imaging in PBS.

## Acknowledgements

We thank W. Li for discussion and A. Choi and B. Unger for comments on the manuscript. This work was supported by the National Institute of General Medical Sciences of the National Institutes of Health (DP2GM132681), the Packard Fellowships for Science and Engineering, and the Bakar Fellows Award, to K.X. K.X. is a Chan Zuckerberg Biohub investigator.

## Author Contributions

K. X. conceived the research. K. C. designed and conducted the experiments. All authors contributed to experimental designs, data analysis, and paper writing.

## References

1. Lichtman, J. W. & Conchello, J. A. Fluorescence microscopy. Nat. Methods 2, 910–919 (2005).

2. Peng, X., Huang, X., Du, K., Liu, H. & Chen, L. High spatiotemporal resolution and low photo-toxicity fluorescence imaging in live cells and in vivo. Biochem. Soc. Trans. 47, 1635–1650 (2019).

3. Tamura, T. & Hamachi, I. Recent progress in design of protein-based fluorescent biosensors and their cellular applications. ACS Chem. Biol. 9, 2708–2717 (2014).

4. Pietraszewska-Bogiel, A. & Gadella, T. W. J. FRET microscopy: from principle to routine technology in cell biology. J. Microsc. 241, 111–118 (2011).

5. Greenwald, E. C., Mehta, S. & Zhang, J. Genetically encoded fluorescent biosensors illuminate the spatiotemporal regulation of signaling networks. Chem. Rev. 118, 11707–11794 (2018).

6. Algar, W. R., Hildebrandt, N., Vogel, S. S. & Medintz, I. L. FRET as a biomolecular research tool - understanding its potential while avoiding pitfalls. Nat. Methods 16, 815–829 (2019).

7. Zimmermann, T., Rietdorf, J. & Pepperkok, R. Spectral imaging and its applications in live cell microscopy.FEBS Lett. 546, 87–92 (2003).

8. Garini, Y., Young, I. T. & McNamara, G. Spectral imaging: Principles and applications. Cytom. Part A 69A, 735–747 (2006).

9. Gao, L. & Smith, R. T. Optical hyperspectral imaging in microscopy and spectroscopy - a review of data acquisition. J. Biophotonics 8, 441–456 (2015).

10. Elliott, A. D. et al. Real-time hyperspectral fluorescence imaging of pancreatic β-cell dynamics with the image mapping spectrometer. J. Cell Sci. 125, 4833–4840 (2012).

11. Wang, Y., Yang, B., Feng, S., Pessino, V. & Huang, B. Multicolor fluorescent imaging by space-constrained computational spectral imaging. Opt. Express 27, 5393–5402 (2019).

12. Zhang, Z., Kenny, S. J., Hauser, M., Li, W. & Xu, K. Ultrahigh-throughput single-molecule spectroscopy and spectrally resolved super-resolution microscopy. Nat. Methods 12, 935–938 (2015).

13. Song, K. H., Zhang, Y., Brenner, B., Sun, C. & Zhang, H. F. Symmetrically dispersed spectroscopic single-molecule localization microscopy. Light Sci. Appl. 9, 92 (2020).

14. Gat, N. Imaging spectroscopy using tunable filters: A review. Proc. SPIE 4056, 50–64 (2000).

15. Favreau, P. et al. Thin-film tunable filters for hyperspectral fluorescence microscopy. J. Biomed. Opt. 19, 011017 (2014).

16. Wachman, E. S., Niu, W. H. & Farkas, D. L. AOTF microscope for imaging with increased speed and spectral versatility. Biophys. J. 73, 1215–1222 (1997).

17. Bei, L., Dennis, G. I., Miller, H. M., Spaine, T. W. & Carnahan, J. W. Acousto-optic tunable filters: fundamentals and applications as applied to chemical analysis techniques. Prog. Quantum Electron. 28, 67–87 (2004).

18. Frank, J. H. et al. A white light confocal microscope for spectrally resolved multidimensional imaging. J. Microsc. 227, 203–215 (2007).

19. Owen, D. M. et al. Excitation-resolved hyperspectral fluorescence lifetime imaging using a UV-extended supercontinuum source. Opt. Lett. 32, 3408–3410 (2007).

20. Favreau, P. F. et al. Excitation-scanning hyperspectral imaging microscope. J. Biomed. Opt. 19, 046010 (2014).

21. Valm, A. M. et al. Applying systems-level spectral imaging and analysis to reveal the organelle interactome. Nature 546, 162–167 (2017).

22. Lewis, S. C., Uchiyama, L. F. & Nunnari, J. ER-mitochondria contacts couple mtDNA synthesis with mitochondrial division in human cells. Science 353, aaf5549 (2016).

23. Wong, Y. C., Ysselstein, D. & Krainc, D. Mitochondria-lysosome contacts regulate mitochondrial fission via RAB7 GTP hydrolysis. Nature 554, 382–386 (2018).

24. Qin, J. et al. ER-mitochondria contacts promote mtDNA nucleoids active transportation via mitochondrial dynamic tubulation. Nat. Commun. 11, 4471 (2020).

25. Olzmann, J. A. & Carvalho, P. Dynamics and functions of lipid droplets. Nat. Rev. Mol. Cell Biol. 20, 137–155 (2019).

26. Tantama, M., Hung, Y. P. & Yellen, G. Imaging intracellular pH in live cells with a genetically encoded red fluorescent protein sensor. J. Am. Chem. Soc. 133, 10034–10037 (2011).

27. Rosselin, M., Santo-Domingo, J., Bermont, F., Giacomello, M. & Demaurex, N. L-OPA1 regulates mitoflash biogenesis independently from membrane fusion. EMBO Rep. 18, 451–463 (2017).

28. Llopis, J., McCaffery, J. M., Miyawaki, A., Farquhar, M. G. & Tsien, R. Y. Measurement of cytosolic, mitochondrial, and Golgi pH in single living cells with green fluorescent proteins. Proc. Natl. Acad. Sci. U. S. A. 95, 6803–6808 (1998).

29. Stryer, L. Fluorescence spectroscopy of proteins. Science 162, 526–533 (1968).

30. Lam, A. J. et al. Improving FRET dynamic range with bright green and red fluorescent proteins. Nat. Methods 9, 1005–1012 (2012).

31. Boersma, A. J., Zuhorn, I. S. & Poolman, B. A sensor for quantification of macromolecular crowding in living cells. Nat. Methods 12, 227–229 (2015).

32. Sukenik, S., Salm, M., Wang, Y. H. & Gruebele, M. In-cell titration of small solutes controls protein stability and aggregation. J. Am. Chem. Soc. 140, 10497–10503 (2018).

33. Sinha, B. et al. Cells respond to mechanical stress by rapid disassembly of caveolae. Cell 144, 402–413 (2011).

34. Hoffmann, E. K., Lambert, I. H. & Pedersen, S. F. Physiology of cell volume regulation in vertebrates. Physiol. Rev. 89, 193–277 (2009).

35. Youle, R. J. & Narendra, D. P. Mechanisms of mitophagy. Nat. Rev. Mol. Cell Biol. 12, 9–14 (2011).

36. Jin, S. M. & Youle, R. J. PINK1-and Parkin-mediated mitophagy at a glance. J. Cell Sci. 125, 795–799 (2012).

37. Katayama, H., Kogure, T., Mizushima, N., Yoshimori, T. & Miyawaki, A. A sensitive and quantitative technique for detecting autophagic events based on lysosomal delivery. Chem. Biol. 18, 1042–1052 (2011).

38. Katayama, H. et al. Visualizing and modulating mitophagy for therapeutic studies of neurodegeneration. Cell 181, 1176–1187 (2020).

39. Wei, L. et al. Super-multiplex vibrational imaging. Nature 544, 465–470 (2017).

40. Niehorster, T. et al. Multi-target spectrally resolved fluorescence lifetime imaging microscopy. Nat. Methods 13, 257–262 (2016).

41. Power, R. M. & Huisken, J. A guide to light-sheet fluorescence microscopy for multiscale imaging. Nat. Methods 14, 360–373 (2017).

42. Wu, Y. & Shroff, H. Faster, sharper, and deeper: structured illumination microscopy for biological imaging. Nat. Methods 15, 1011–1019 (2018).

43. Poudel, C. & Kaminski, C. F. Supercontinuum radiation in fluorescence microscopy and biomedical imaging applications. J. Opt. Soc. Am. B-Opt. Phys. 36, A139–A153 (2019).

