## Supplemental figures for "Excitation spectral microscopy for highly multiplexed fluorescence imaging and quantitative biosensing"

### SUPPLEMENTAL INFORMATION

#### Supplementary Figures

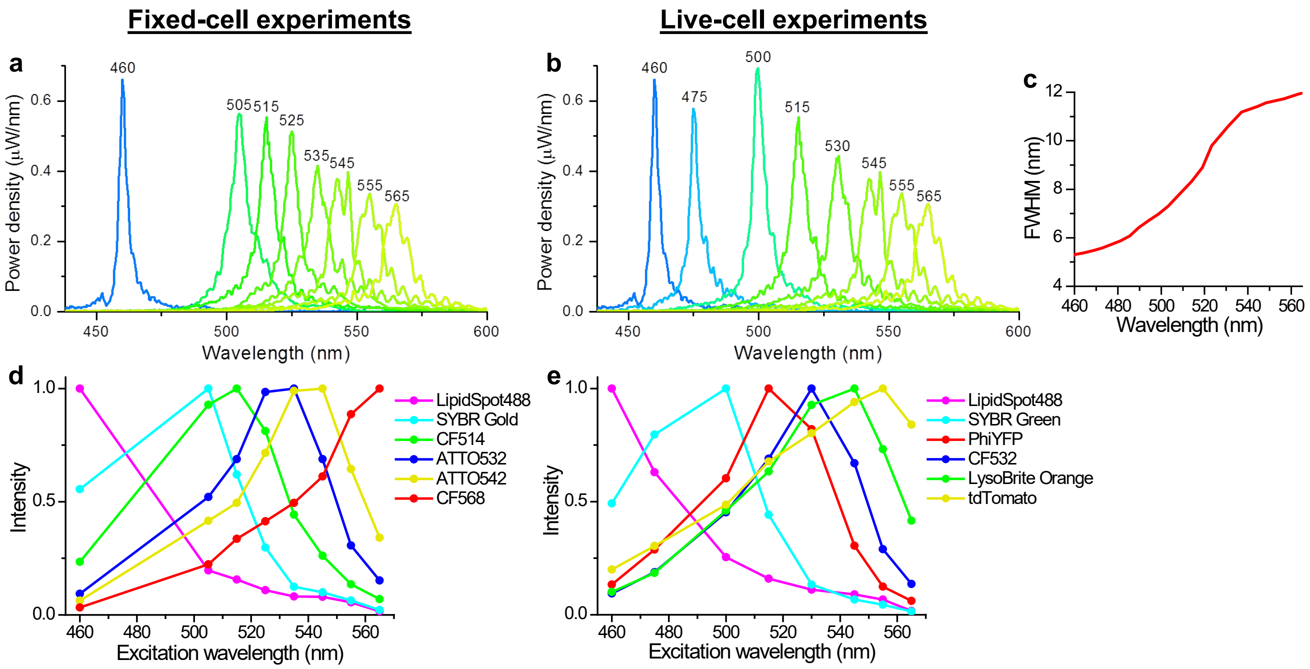


**Supplementary Fig. S1. | Excitation spectral profiles and their expected effects on the recorded spectra.** (a, b) Measured AOTF output (microscope input) spectral profiles for the 8 excitation wavelengths used for the 10 fps fixed-cell (a) and live-cell (b) experiments. (c) Full width at half maximum (FWHM) of the AOTF output as a function of the central wavelength. (d, e) The expected spectral responses of different fluorophores at the 8 excitation wavelengths for our 6-target fixed-cell (d) and 6-target live-cell (e) experiments, respectively, obtained by multiplying the power-normalized spectral profiles in (a, b) with the literature absorption spectra of the fluorophores used in the two experiments. The non-ideal AOTF spectral profiles thus distort the original spectra. Yet, as long as the resultant spectral responses of different fluorophores are still distinct, robust linear unmixing may still be achieved (Supplementary Figs. S2 and S3 below). The above-predicted spectral responses of different fluorophores are in general agreement with our experimental results (Figs. 1c and 2c); small deviations are attributed to the possible differences between the literature absorption spectra and the actual excitation spectra of the fluorophores.

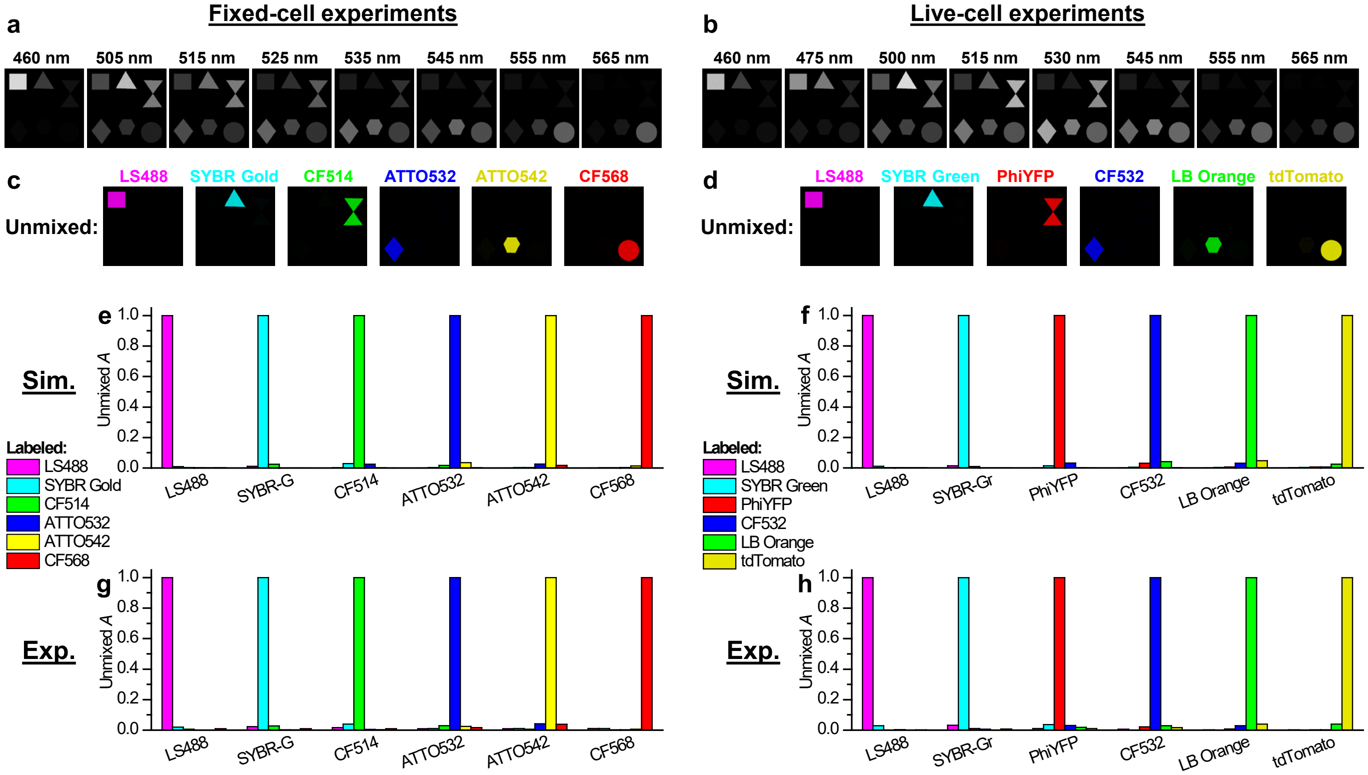


**Supplementary Fig. S2. | A comparison of the simulated and experimental unmixing results.** (a, b) Simulated 8-wavelength images for the fluorophores used in our fixed-cell (a) and live-cell (b) experiments: each of the 6 fluorophores in either experiment was simulated based on the expected excitation spectra in Supplementary Figs. S1d (for a) and S1e (for b) to occupy a different region of the view, and then an uncorrelated Gaussian noise of a standard deviation 5% of the signal was added to every pixel at each wavelength. (c, d) Unmixing of the 8-wavelength images in (a, b) into the 6 fluorophore channels. (e, f) The unmixed abundancy values in different fluorophore channels for the different simulated regions occupied by each of the fluorophores. (g, h) Corresponding experimental unmixing results for samples singly labeled by each of the fluorophores.

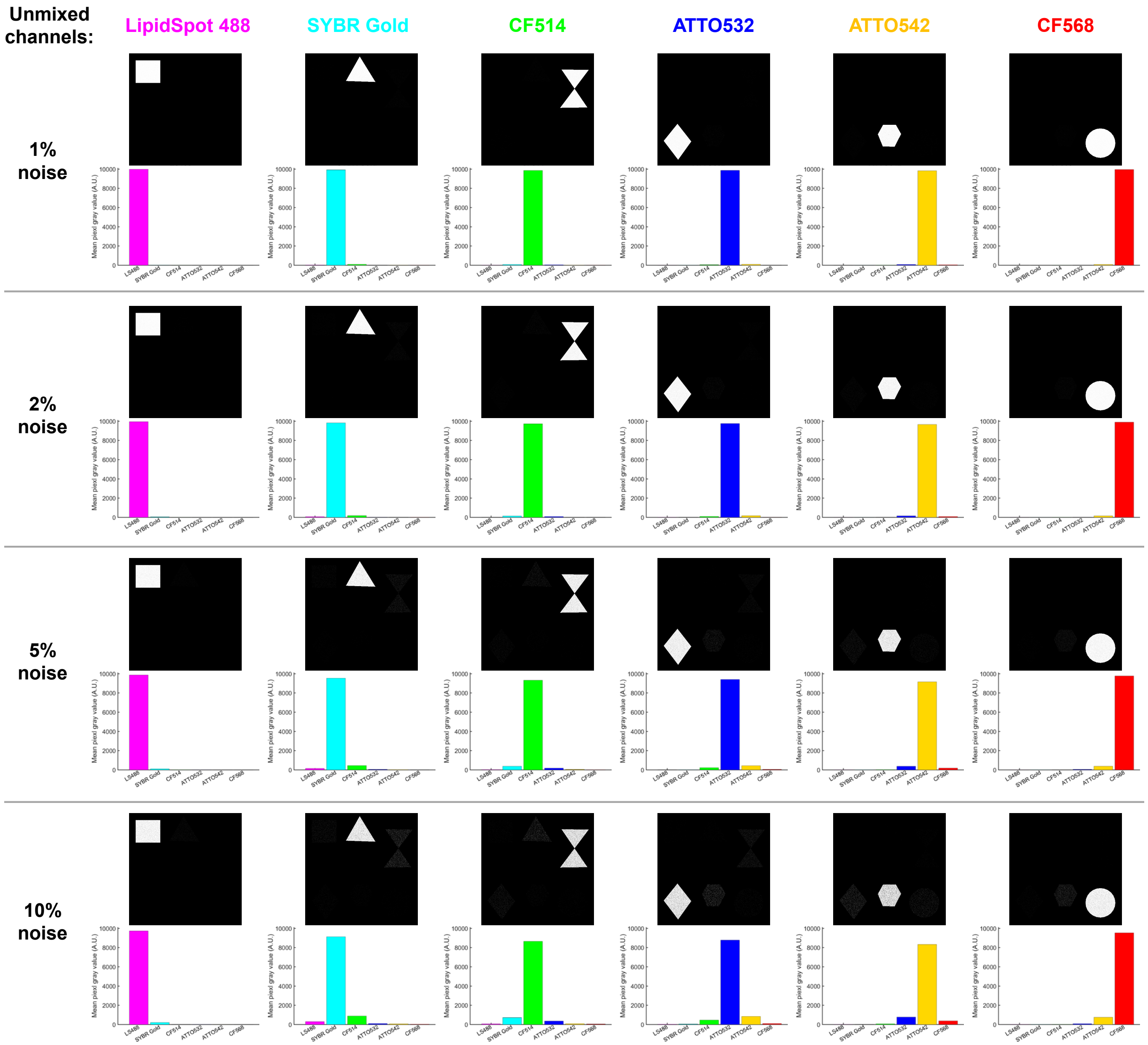


**Supplementary Fig. S3. | Unmixing results for simulated data based on the experimental excitation spectra, at different noise levels.** In this simulation, each of the 6 fluorophores in our fixed-cell experiments was simulated based on the experimental excitation spectra (Fig. 1c) to occupy a different region of the view, and then uncorrelated Gaussian noises of standard deviations of 1%, 2%, 5%, and 10% of the signal were added to every pixel at each wavelength. For each noise level, the top row presents the unmixed images in each channel, and the bottom row gives the average intensities observed in the corresponding channel for each labeled region.

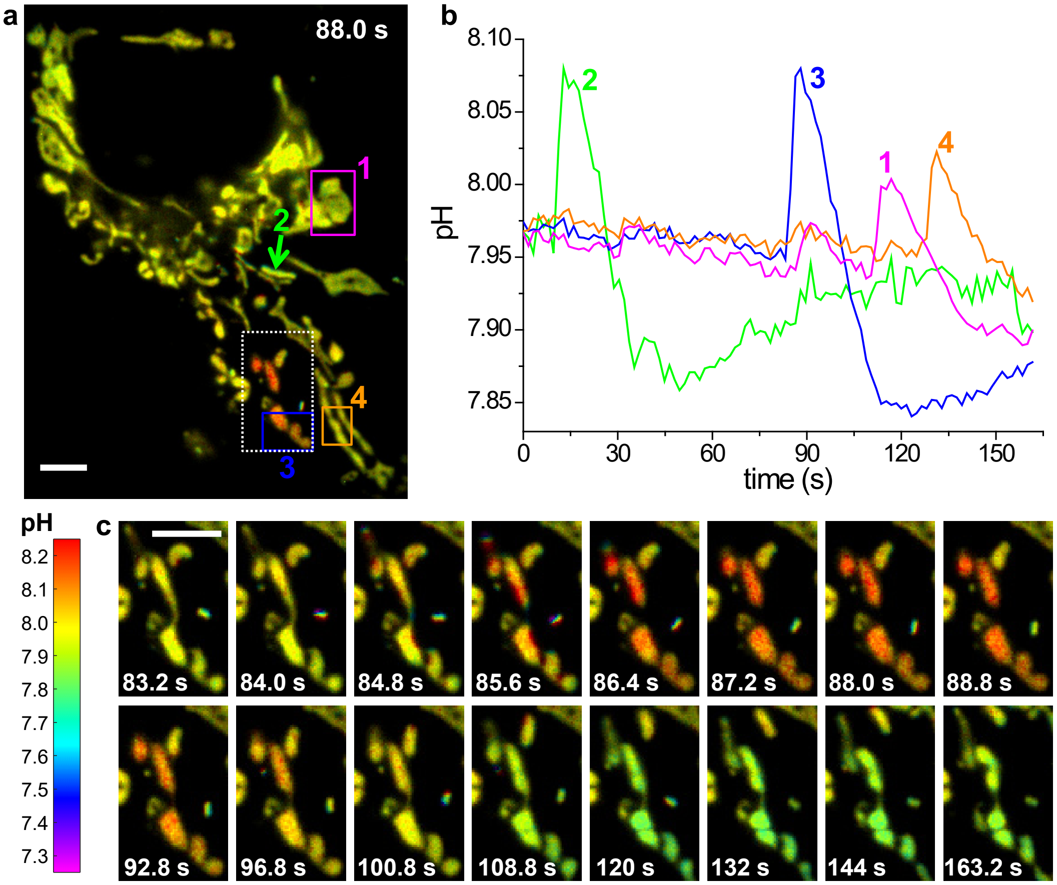


**Supplementary Fig. S4. | Another example for the typical dynamics of the mitochondrial matrix pH in another live COS-7 cell.** (a) Color-coded Mito-pHRed absolute pH map at the time point of 88.0 s, presented on the same color scale as Fig. 3. (b) pH time traces for the four regions/mitochondria marked in (a). (c) Time sequences for the white-box regions in (a). The top row presents consecutive pH images at 0.8 s time spacing. The bottom row presents pH images at larger time separations for the slower dynamics of pH recovery. Experiment was performed with 8-wavelength excitation at 10 fps (0.8 s acquisition time for each spectral image). Scale bars: 5 μm (a, c). See Supplementary Video 7 for a movie of (a).

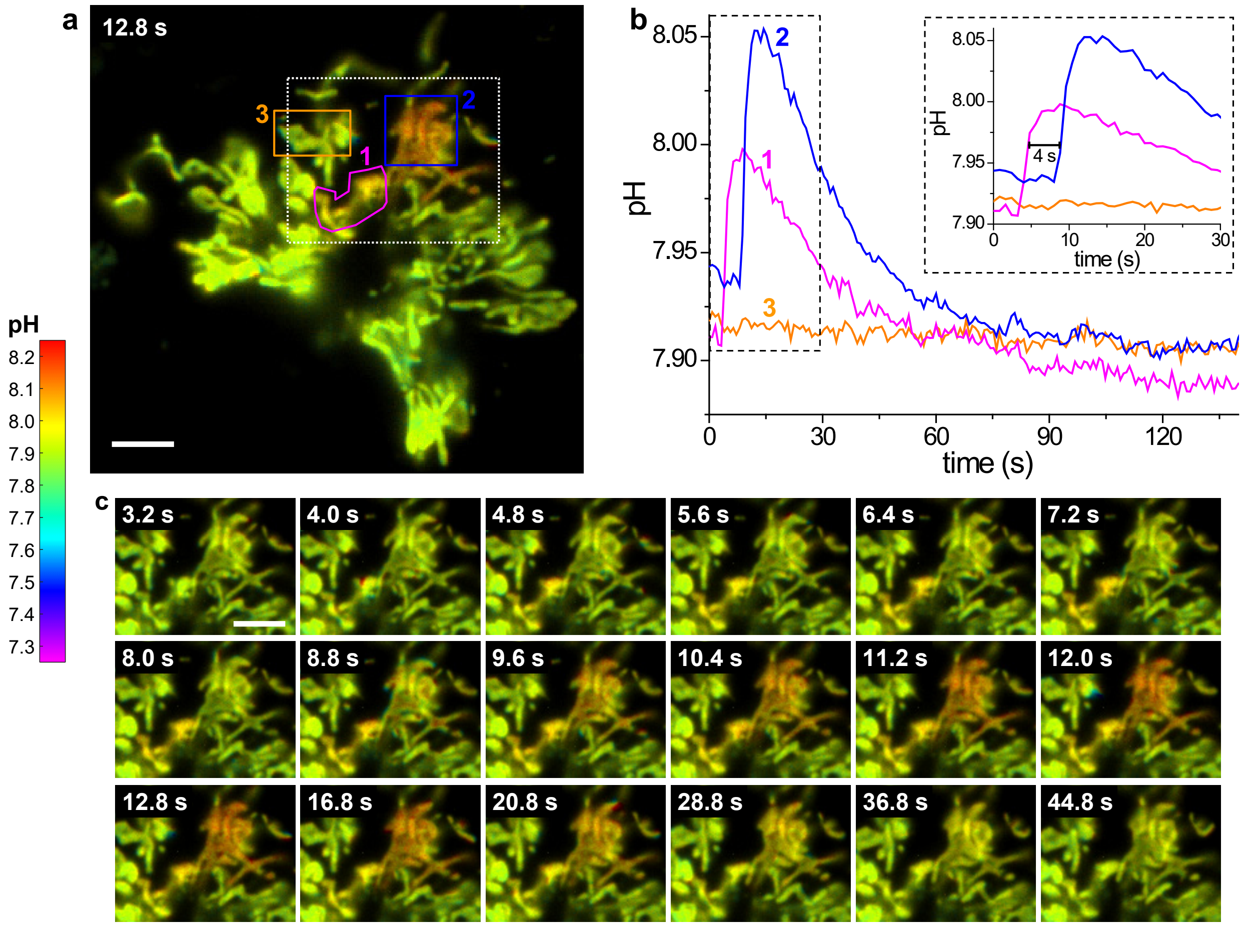


**Supplementary Fig. S5. | Mitochondrial matrix pH dynamics in another live COS-7 cell that showed tandem pH jumps for two neighboring regions.** (a) Color-coded Mito-pHRed absolute pH map at the time point of 12.8 s, presented on the same color scale as Fig. 3. (b) pH time traces for the 3 regions marked in (a). Inset: zoom-in of the first 30 s. The pH jump of Region 2 started 4 s later after Region 1, at which time the pH in Region 1 already peaked. Meanwhile, the pH of Region 3 remained unchanged over the observation timeframe. (c) Time sequences for the white-boxed regions in (a). The top two rows give consecutive pH images recorded at 0.8 s time spacing. The bottom row is presented at larger time separations for the slower dynamics of pH recovery. Experiment was performed with 8-wavelength excitation at 10 fps (0.8 s acquisition time for each spectral image). Scale bars: 5 μm (a, c).

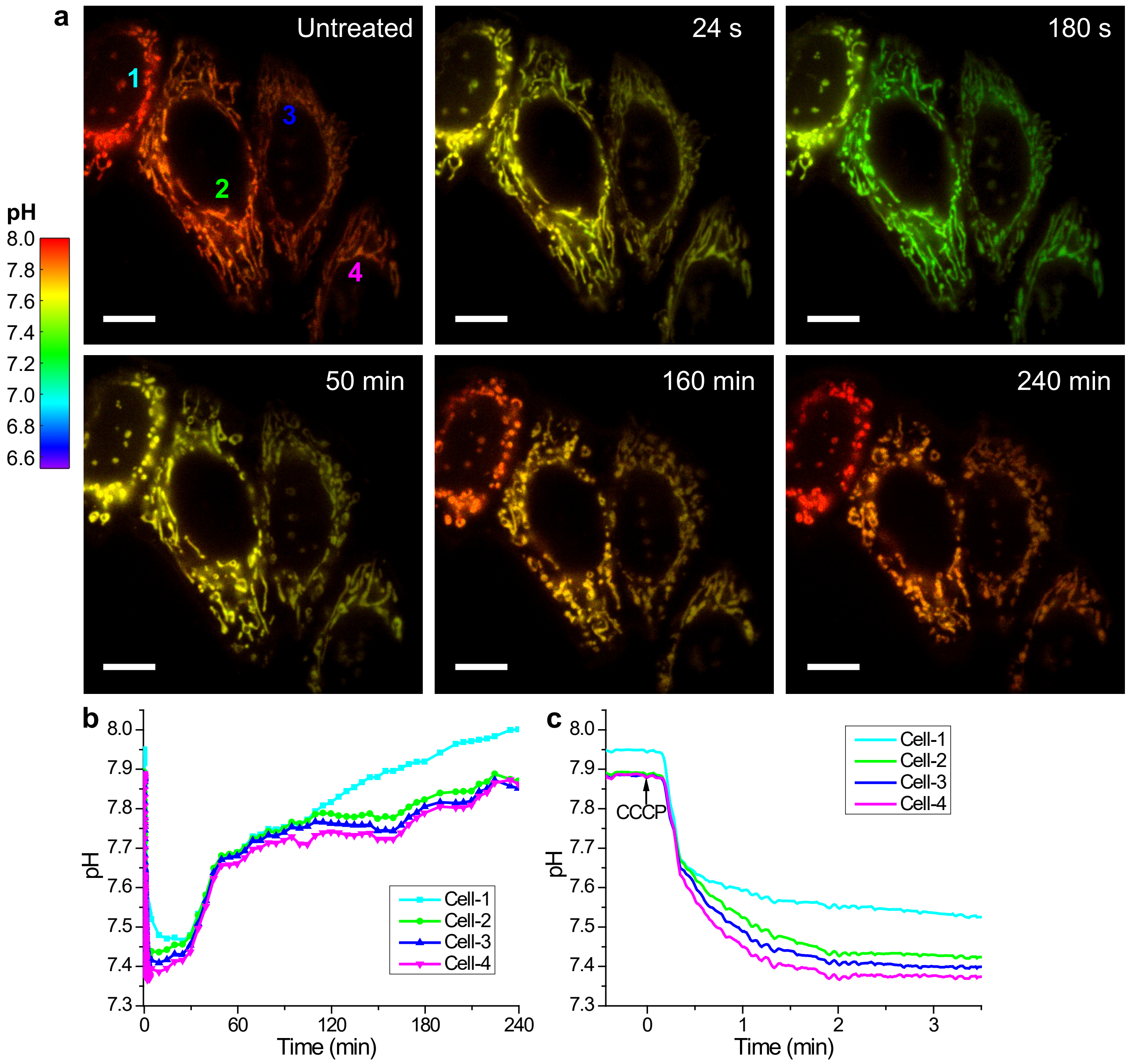


**Supplementary Fig. S6. | Long-term evolution of the mitochondrial matrix pH in live HeLa cells after CCCP treatment.** (a) Color-coded Mito-pHRed absolute pH maps for HeLa cells not expressing Parkin, before and after the application of 20 μM CCCP at different time points. To facilitate a direct comparison with the mitophagy data, the same pH color scale as Fig. 5 is used, different from the color scale used in Fig. 3. (b) Averaged pH for the mitochondria in the four cells marked in (a), as a function of time. (c) Zoom-in of the initial fast pH drop. Experiments were performed with 8-wavelength excitation at 10 fps (0.8 s acquisition time for each spectral image), with continuous recording for $t < 3.5$ min and intermittent recording every 5 min for the later time points. Scale bars: 10 μm.

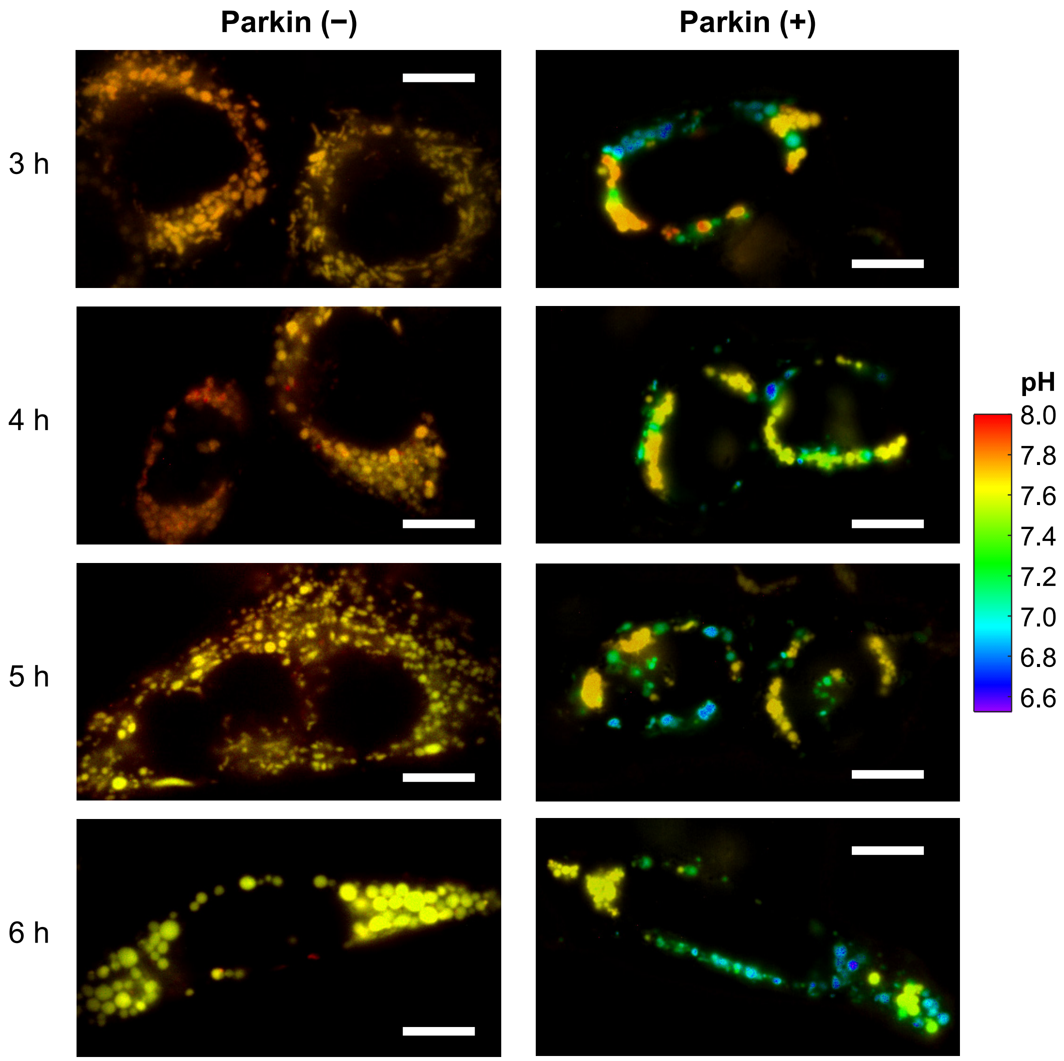


**Supplementary Fig. S7. | Representative Mito-pHRed pH maps for live HeLa cells not expressing (left) and expressing (right) fluorescently untagged Parkin, after the application of 20 μM CCCP for different times.** Experiments were performed with 8-wavelength excitation at 10 fps (0.8 s acquisition time for each spectral image). Scale bars: 10 μm.

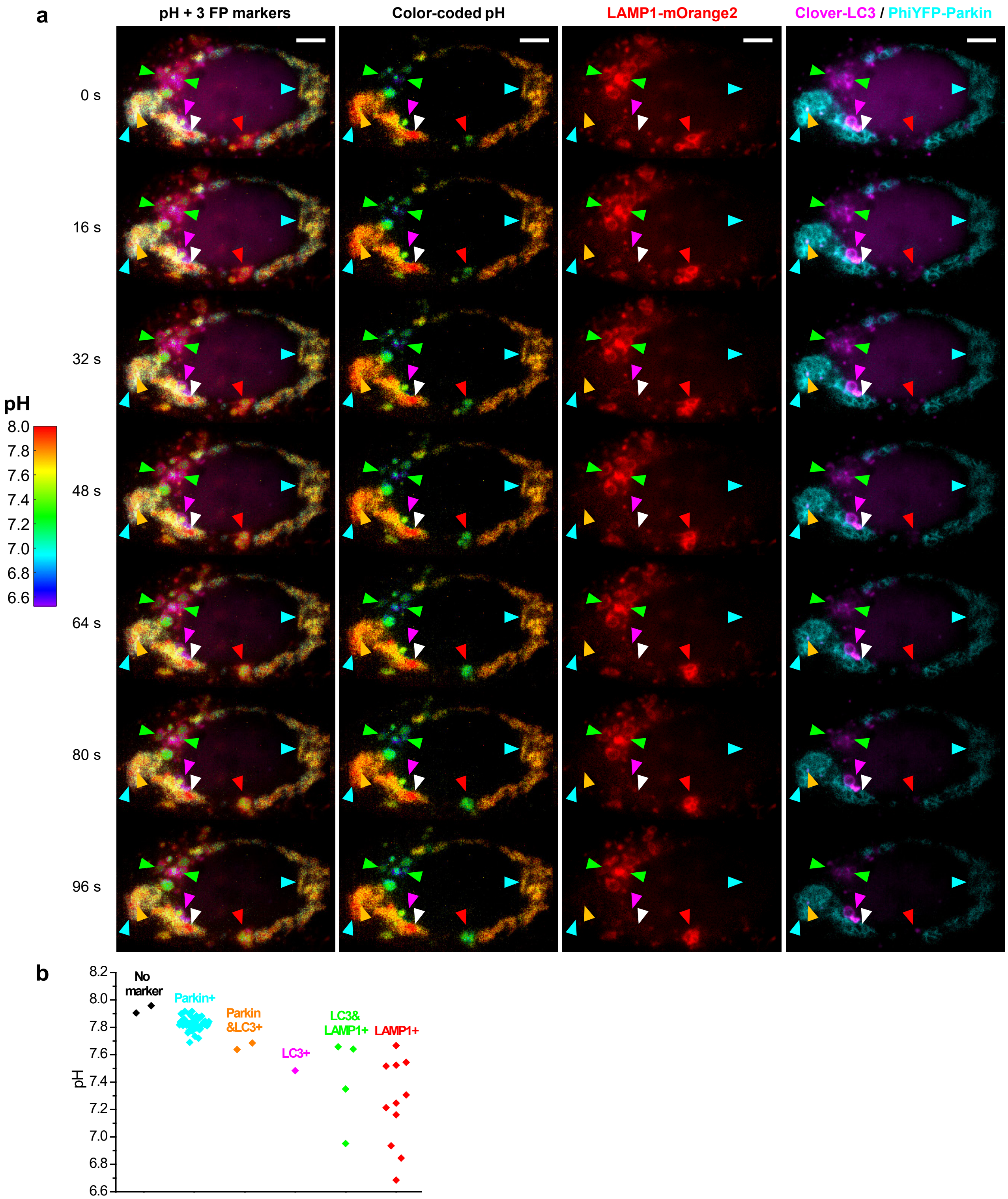


**Supplementary Fig. S8. | Concurrent absolute pH imaging of the mitochondrial matrix with three FP markers in the Parkin-mediated mitophagy pathway, with a different permutation of FPs.** (a) Image series, with overlaid and separately presented images of color-coded Mito-pHRed absolute pH map, LAMP1-mOrange2, and Clover-LC3/PhiYFP-Parkin, for a Parkin-expressing live HeLa cell after the application of 20 μM CCCP for 4 h. Arrowheads point to representative vesicles that are: not labeled by any markers (white), labeled by Parkin only (cyan), labeled by both Parkin and LC3 (orange), labeled by LC3 only (magenta), labeled by both LC3 and LAMP1 (green), and labeled by LAMP1 only (red). (b) Distribution of the Mito-pHRed-quantified matrix pH for individual mitochondria labeled by the different markers. Scale bars: 5 μm (a). Experiment was performed with 8-wavelength excitation at 10 fps, and images were generated from averages of 5 excitation cycles for presenting at a time spacing of 4 s. See Supplementary Video 10 for the full image series.

##### Supplementary Video Captions

**Supplementary Video 1.** Movie of the 6-color live-cell data shown in Fig. 2b. "0" time corresponds to the time point of Fig. 2b. Scale bar: 10 μm.

**Supplementary Video 2.** Movie of a section of the same 6-color live-cell data (Left), compared to the same data after removing the cell-membrane and ER channels (Right) for a clearer presentation of the other 4 channels. Scale bar: 2 μm.

**Supplementary Video 3.** Movie for the fast 4-color live-cell data of Fig. 2e. "0" time corresponds to the time point of the left panel of Fig. 2e. Scale bar: 5 μm.

**Supplementary Video 4.** Movie for the ER-mediated fusion of two lipid droplets shown in Fig. 2hi. Scale bar: 1 μm.

**Supplementary Video 5.** Movie for the Mito-pHRed absolute pH map of Fig. 3e, for a COS-7 cell that was treated with CCCP. "0" time corresponds to the time point of CCCP addition. See color scale for pH in Fig. 3. Scale bar: 10 μm.

**Supplementary Video 6.** Movie for the Mito-pHRed absolute pH map of Fig. 3g. See color scale for pH in Fig. 3. Scale bar: 5 μm.

**Supplementary Video 7.** Movie for the Mito-pHRed absolute pH map of Supplementary Fig. S4. See pH color scale in Supplementary Fig. S4. Scale bar: 5 μm.

**Supplementary Video 8.** Movie for the color-coded FRET map of the crowding sensor in Fig. 4e, for two COS-7 cells that underwent 150% hypertonic treatment. "0" time corresponds to the time point of sorbitol addition. See color scale for FRET efficiency in Fig. 4. Scale bar: 10 μm.

**Supplementary Video 9.** Movie for the color-coded FRET map of the crowding sensor in Fig. 4f, for two COS-7 cells that underwent 50% hypotonic treatment. "0" time corresponds to the time point of water addition. See color scale for FRET efficiency in Fig. 4. Scale bar: 10 μm.

**Supplementary Video 10.** Video for the concurrent absolute pH imaging with 3 additional FPs shown in Supplementary Fig. S8, shown as an overlaid image and separated absolute pH colormap and FP channels. See pH color scale and color coding for the 3 FPs in Supplementary Fig. S8. Arrows point to LAMP1-marked mitochondria exhibiting fast dynamics. Scale bar: 5 μm.
